# Loss of multiple enzyme activities due to the human genetic variation P284T in NADPH cytochrome P450 oxidoreductase

**DOI:** 10.1101/643825

**Authors:** Shaheena Parween, Maria Natalia Rojas Velazquez, Sameer S. Udhane, Norio Kagawa, Amit V. Pandey

## Abstract

Cytochromes P450 located in the endoplasmic reticulum require NADPH cytochrome P450 oxidoreductase (POR) for their catalytic activities. Mutations in POR cause multiple disorders in humans related to the biosynthesis of steroid hormones and also affect drug-metabolizing cytochrome P450 activities. Here we are reporting the effects of a POR genetic variant P284T which is located in the hinge region of POR that is necessary for the flexibility of domain movements. Human wild-type and P284T mutant of POR, as well as cytochrome P450 proteins, were expressed in bacteria, purified and then reconstituted in liposomes for enzyme kinetic assays. Quality of POR proteins was checked by cytochrome c, ferricyanide and tetrazolium dye reduction assay and measurements flavin content. We found that for the P284T variant of POR the cytochrome c reduction activity was reduced to 47% of the WT and MTT reduction was reduced to only 15% of the WT. No impact on ferricyanide reduction activity was observed, but a severe loss of CYP19A1 (aromatase) activity was observed (9% of WT). In the assays of drug metabolizing cytochrome P450 enzymes, the P284T variant of POR showed 26% activity for CYP2C9, 44% activity for CYP2C19, 23% activity for CYP3A4 and 44% activity in CYP3A5 assays compared to the WT POR. These results indicate a severe effect on several cytochrome P450 activities due to the P284T variation in POR which suggests a negative impact on both the steroid as well as drug metabolism in the individuals carrying this variation.

## 1 Introduction

The NADPH cytochrome P450 reductase (POR, NCBI# NP_000932, UniProt# P16435) is the obligate electron donor for all cytochromes P450 proteins located in the endoplasmic reticulum (Pandey and Flück, 2013;Flück and Pandey, 2019). Cytochrome P450 proteins are responsible for the metabolism of numerous endogenous and exogenous chemicals, including most of the drugs, steroids and other xenobiotics (Schuster and Bernhardt, 2007;Omura, 2010;Zanger and Schwab, 2013). There are two different types of cytochrome P450 proteins in mammals (Omura, 2010;McLean et al., 2015). The type 1 P450s are found in the mitochondria use adrenodoxin/adrenodoxin reductase system as their redox partner (Omura, 2006;Zalewski et al., 2016). The type 2 cytochrome P450s are located in the endoplasmic reticulum and use a single protein, POR, as their redox partner (Lu et al., 1969;Masters, 2005). POR contains both the flavin mononucleotide (FMN) and the flavin adenine dinucleotide (FAD) which are bound to distinct domains and supplies electrons to cytochromes P450 and other proteins via protein-protein interactions (Iyanagi and Mason, 1973;Iyanagi et al., 1974;Matsubara et al., 1976;Vermilion and Coon, 1978;Oprian and Coon, 1980) (**Figure 1**). The three-dimensional structure of the FMN binding domain of human POR has been solved by x-ray crystallography (Zhao et al., 1999), and more recently many structures of soluble POR proteins containing the NADPH, FAD, and FMN binding domains but lacking the membrane-bound amino terminus sequence has been determined by several groups (Wang et al., 1997;Aigrain et al., 2009;Xia et al., 2011a;Xia et al., 2011b).

**Figure 1:**
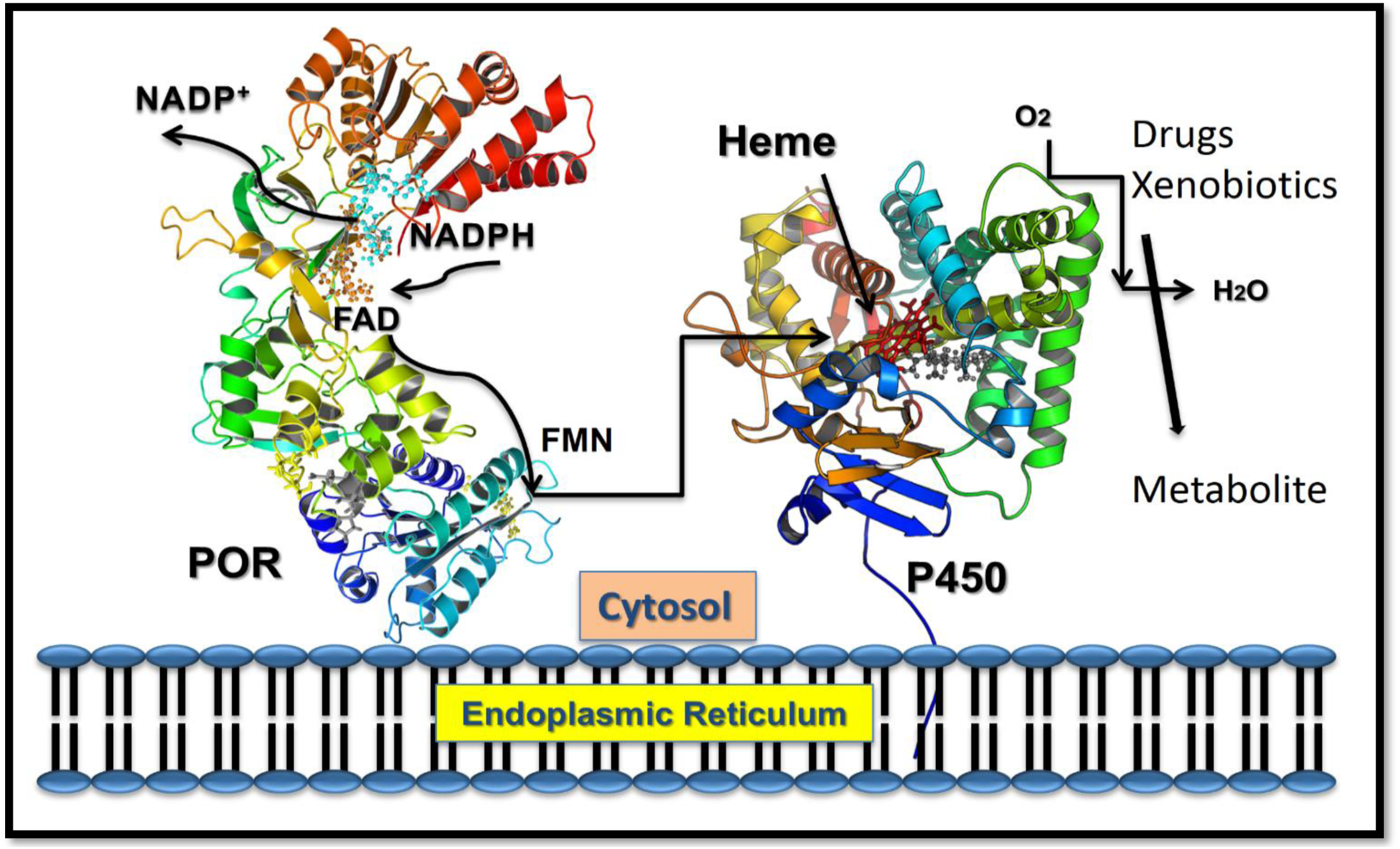
POR as electron transfer partner to cytochromes P450 proteins. Transfer of electrons from NADPH to redox partners of POR. Co-factor NADPH binds to the POR located into the endoplasmic reticulum and donates electrons which are received by FAD. Electron transfer to FAD causes a conformational change in POR that brings together the FAD, and FMN domains and electrons are then transferred from FAD to FMN. The FMN domain of POR interacts with the P450s and other redox partners and completes the final step of electron transfer. From Pandey, A. V. and Sproll, P (2014). “Pharmacogenomics of human P450 oxidoreductase.” *Frontiers in Pharmacology* **5**, 103.

Human POR deficiency (PORD, OMIM: 613537 and 201750) is a form of congenital adrenal hyperplasia, initially described in patients with altered steroidogenesis (Flück et al., 2004;Miller et al., 2004;Flück et al., 2007;Flück and Pandey, 2011;Flück and Pandey, 2013;Pandey and Flück, 2013;Fukami and Ogata, 2014;Pandey and Sproll, 2014;Flück and Pandey, 2019) followed by several reports with a broad spectrum of disorders (Adachi et al., 2004;Arlt et al., 2004;Pandey and Flück, 2013;Flück and Pandey, 2014;Pandey and Sproll, 2014;Oldani et al., 2015;Bonamichi et al., 2016;Nakanishi et al., 2016;Burkhard et al., 2017;Flück and Pandey, 2019). Congenital adrenal hyperplasia is an inborn error of adrenal steroid biosynthesis affecting the production of glucocorticoids (Miller and Fluck, 2014;Fluck, 2017). Sequencing of the POR gene in patients with symptoms of mixed oxidase deficiency revealed mutations in POR linked to disorders of steroid biosynthesis (Flück et al., 2004;Miller et al., 2004;Pandey et al., 2004). Afterward, several different laboratories reported mutations in POR in patients with steroid biosynthesis disorders and bone malformation syndromes (Adachi et al., 2004;Arlt et al., 2004;Fukami et al., 2005;Pandey and Flück, 2013;Riddick et al., 2013). While the initial studies on mutations in POR were on the metabolism of steroid hormones, effects of POR on other redox partners including heme oxygenase and drug metabolizing cytochromes P450 have been studied in later reports (Marohnic et al., 2006;Agrawal et al., 2008;Kranendonk et al., 2008;Agrawal et al., 2010;Flück et al., 2010;Marohnic et al., 2010;Nicolo et al., 2010;Pandey et al., 2010;Marohnic et al., 2011;McCammon et al., 2016;Udhane et al., 2017). A mouse knockout (KO) model of *Por-/-* had been generated before the human mutations in POR were identified. The *Por-/-* was found to be embryonic lethal (Shen et al., 2002;Nicholas et al., 2017), and resembled the KO mouse model of retinoic acid-metabolizing *Cyp26a1-/-* (Gaston-Massuet et al., 2016). Liver specific deletion of POR has been reported to cause lipidosis and increased liver weight (Gu et al., 2003;Wu et al., 2003). Deletion of POR gene in cardiomyosite cells of mouse lead to reduced P450 activity in heart but there was no indication of embryonic lethality or early morbidity and no difference in heart function was observed (Fang et al., 2008).

Large-scale sequencing projects have now identified many variations in the POR gene, in several different human subpopulations, and multiple POR variants have been linked to altered drug metabolism (Gomes et al., 2008;Huang et al., 2008;Gomes et al., 2009;Agrawal et al., 2010;Nicolo et al., 2010;Pandey et al., 2010). Since POR is necessary for several biochemical reactions of steroid biosynthesis that are catalyzed by cytochrome P450 proteins, a complex disorder is caused by mutations in POR. Several variants in the POR gene have been found in patients and non-clinical samples, have been tested for enzymatic activities (Adachi et al., 2004;Arlt et al., 2004;Fukami et al., 2005;Fukami et al., 2006;Homma et al., 2006;Pandey, 2006;Huang et al., 2008;Sim et al., 2009).

While the mutations in POR may be located in all regions of the protein, based on analysis of previously identified mutations, some definite patterns have emerged. Mutations that are located near the co-factor binding sites (FMN, FAD, and NADPH) generally give rise to a severe form of the disease with the mutations causing severe loss of FMN or FAD showing most significant inhibitory effects on activities of all redox partners. We have previously shown that aromatase (CYP19A1) activity responsible for the metabolism of androgens to form estrogens is more susceptible to changes in NADPH binding site mutations in POR (Pandey et al., 2007;Flück et al., 2011;Flück and Pandey, 2017).

In the current study, we have investigated the effects of POR variant P284T located near the hinge region of POR that is crucial for interactions with redox partner proteins, including P450s, to transfer electrons from NADPH. The POR variant P284T (rs72557937) has been identified in the Yoruba population through the sequencing of DNA obtained from a particular community in Ibadan, Nigeria (https://www.coriell.org/0/Sections/Collections/NHGRI/Yoruba.aspx?PgId=128). Based on information available through the 1000 genome project, the samples were from parent-adult child trios, and all parents in the trios had identified themselves as having four Yoruba grandparents. The P284T variant of POR was found in the African American population in the POR sequencing study carried out by Miller (www.pharmagenomics.unsf.edu/populations.html).

## 2 Materials and methods

### 2.1 Recombinant Expression of POR and Membrane Purification

The human POR WT and variant forms of proteins (NCBI# NP_000932, Uniprot# P16435) were recombinantly expressed in bacteria using the previously described methods and expression constructs (Huang et al., 2005;Pandey et al., 2007;Huang et al., 2008;Nicolo et al., 2010). The protocol for expression of an N-27 form of POR variants and subsequent membrane purification have been described and adopted from our previous publications (Huang et al., 2005;Pandey et al., 2007;Pandey et al., 2010;Parween et al., 2016;Flück and Pandey, 2017). The cDNAs for WT or mutant POR in a pET22b vector were transformed into Escherichia coli BL21(DE3). Single colonies were selected for growth on ampicillin and grown in terrific broth pH 7.4 supplemented with 40 mM FeCl_3_, 4 mM ZnCl_2_, 2 mM CoCl_2_, 2 mM Na_2_MoO_4_, 2 mM CaCl_2_, 2 mM CuCl_2_, 2 mM H_3_BO_3_, 0.5 mg/ml riboflavin, 100 µg/ml carbenicillin at 37 °C to an optical density (OD) 600 nm of 0.6 and temperature was reduced to 25 °C for 16 h. The bacterial cells were collected by centrifugation, washed with PBS and suspended in 100 mM Tris-acetate (pH 7.6), 0.5 M sucrose, and 1 mM EDTA and treated with lysozyme (0.5 mg/ml) and EDTA (0.1mM [pH 8.0]) at 4 °C for 1 h with slow stirring to generate spheroplasts. The spheroplasts were pelleted by centrifugation at 5000×g for 15 min; and suspended in 100 mM potassium phosphate (pH 7.6), 6 mM Magnesium acetate, 0.1 mM DTT, 20% (v/v) glycerol, 0.2 mM PMSF, and 0.1 mM DNase I; and disrupted by sonication. A clear lysate devoid of cellular debris was obtained by centrifugation at 12,000×g for 10 min, and then the membranes were collected by centrifugation at 100,000×g for 60 min at 4 °C. Membranes containing POR were mixed in 50 mM Potassium phosphate buffer (pH 7.8) and 20% (v/v) glycerol and kept at −70 °C. The concentrations of proteins were measured by the RC-DC protein assay method (Protein Assay Dye Reagent, Bio-Rad, Hercules, CA) and POR content in membrane proteins was measured by western blot analysis.

### 2.3 Western Blot Analysis of POR Content in the Bacterial Membranes

For Western blots, 1 μg of POR-WT and POR-P284T bacterial membrane proteins were separated on an SDS-PAGE gel and blotted on to polyvinyl difluoride (PVDF) membranes. Blots were first incubated with a rabbit polyclonal antibody against wild-type human POR from Genscript (Genscript, NJ, USA) at a dilution of 1:1000. We then used a secondary goat anti-rabbit antibody labeled with a phthalocyanine infrared dye (IRDye 700DX, LI-COR Bioscience Inc., NE, USA) at a 1:10000 dilution. Signals were analyzed with the green fluorescent channel (700 nm) on an Odyssey Infrared Imaging System (LI-COR Bioscience Inc., NE, USA), and protein bands were measured using the Odyssey software (LI-COR Bioscience Inc., NE, USA). POR content of each membrane preparation was measured, and all samples were normalized against purified wild-type POR used as a standard. In all experiments described in this report, the normalized amount of POR content was used for the mutants as well as WT POR protein.

### 2.4 Small molecule Reduction Assay by WT and POR-P284T

Assay of cytochrome c reduction by bacterially expressed WT or POR-284T was performed as described previously by measuring the change in absorbance at 550 nm (ε=21.1 cm^-1^ mM^-1^) (Guengerich et al., 2009). In brief, the POR reduction reaction was performed in 96-well plates, in triplicate, with 5 µg membrane preparation containing POR in each well in 100 mM Tris-HCl (pH 7.5), using a microplate reader (Spectramax M2e, Molecular Devices, Sunnyvale, CA). The concentration of NADPH used was 100 μM and concentrations of cytochrome c (1.3–40 μM) were varied for kinetic analysis. The POR reduction reaction was initiated by the addition of NADPH and the change in absorbance at 550 nm was monitored over 6 minutes. Data were fitted based on Michaelis-Menten kinetics (Michaelis and Menten, 1913) using GraphPad Prism (GraphPad Software, La Jolla, CA USA) to determine the Vmax and Km.

The NADPH-dependent MTT [3-(4,5-dimethylthiazol-2-yl)-2,5-diphenyltetrazolium] reduction rate was measured as the rate of increase in absorbance at 610 nm using an extinction coefficient of ε_610_=11 mM-1 cm-1 (Yim et al., 2005). The assay mixture contained 5 μg of bacterial membranes containing POR in 100 mM phosphate buffer (pH 7.6), 100μM NADPH and concentration of MTT varied from 0-500 μM. Similarly, the ferricyanide reduction rate was measured as the rate of decrease in absorbance at 420 nm (ε_420_=1.02 mM-1 cm-1). The concentration of ferricyanide varied over a range from 0 to 500 μM and the reactions were started by adding 100μM NADPH (Marohnic et al., 2010). Activities represent the mean of at least triplicate determinations.

### 2.5 Flavin Content Analysis of WT and Mutant POR

Protein-bound flavin molecules were released by thermal denaturation of POR proteins (Faeder and Siegel, 1973). Flavin content of WT and mutant POR proteins (100 µg/ml) was determined by boiling protein samples at 95 °C for 10 min in the dark, followed by centrifugation at 14000 x g for 10 min to remove coagulated protein. The FMN and FAD ratio was determined by measurement of fluorescence of the supernatant at pH 7.7 and pH 2.6 (excitation at 450 nm, emission at 535 nm) (Faeder and Siegel, 1973).

### 2.6 Expression and Purification of human CYP19A1

The vector for bacterial expression of human CYP19A1was transformed in E. coli BL21(DE3) cells, and the recombinant protein was expressed and purified following previously published protocols (Kagawa et al., 2003;Kagawa, 2011), with slight modification**s**. Briefly, a single transformed colony was selected for protein expression at 25 °C. After 4 h of incubation, 1 mM δ-aminolevulinic acid (a heme precursor) and 4 mg/ml arabinose (for induction of molecular chaperones GroEL/GroES) were added to the culture and further incubated for 20 h. Cells were harvested, and spheroplasts were prepared with 0.2 mg/ml lysozyme in 50 mM Tris-Acetate (pH 7.6), 250 mM sucrose and 0.5 mM EDTA at stored at −80 °C. For protein purification, spheroplasts were lysed using 10XCellLytic B (Sigma-Aldrich) in buffer containing 100mM potassium phosphate (pH 7.4), 500 mM sodium acetate, 0.1 mM EDTA, 0.1 mM DTT, 20% glycerol, and 1 mM PMSF. The cell lysate was centrifuged, and the supernatants were pooled for purification by Ni^2+^ affinity chromatography. Purification was performed at 4 °C and protein concentration after the dialysis was determined by DC protein assay (Protein Assay Dye Reagent, Bio-Rad, Hercules, CA) using BSA as standard.

### 2.7 Assay of cytochrome P450 CYP19A1 in Reconstituted Liposome System

Purified recombinant CYP19A1 using the bacterial expression system was used to test the effect of POR-284T to support the aromatase activity of CYP19A1. Standard tritiated water release assay for the CYP19A1 activity was performed in a reconstituted liposome system using androstenedione as a substrate. Bacterial membranes containing POR and purified CYP19A1 were reconstituted into DLPC-DLPG liposomes. The liposomes were prepared previously (Udhane et al., 2017), and aromatase activity was measured by the tritiated water release assay originally described by Lephart and Simpson (Lephart and Simpson, 1991) using androstenedione as the substrate. Reaction mixture consisted of 100 pmol of CYP19A1, 400 pmol of POR, 100 mM NaCl and ^3^H labeled androstenedione ([1β-^3^H(N)]-andros-tene-3,17-dione; ∼20,000 cpm) in 100 mM potassium phosphate buffer (pH 7.4). Different concentrations (10–1000 nM) of androstenedione were used for kinetic analysis. The catalytic reaction was initiated by the addition of 1 mM NADPH, and the reaction tube was incubated for 1 h under shaking. Data were fitted based on Michaelis-Menten kinetics using GraphPad Prism (GraphPad Software, La Jolla, CA USA).

### 2.8 Assay of cytochrome P450 CYP2C9 Activity in Reconstituted Liposome System

The activity of CYP2C9 promoted by WT or mutant POR was tested using the fluorogenic substrate BOMCC (Invitrogen Corp, Carlsbad, CA, United States). The purified CYP2C9 (CYPEX, Dundee, Scotland, United Kingdom) was used to test the activities of the POR variants using 20 µM BOMCC as substrate. In vitro CYP2C9 assays were performed using a reconstituted liposome system consisting of WT/mutant POR, CYP2C9 and cytochrome b5 at a ratio of 5:1:1. The final assay mixture consisted of 5 µg DLPC (1,2-Dilauroyl-sn-glycero-3-phosphocholine) and proteins (1 µM POR: 200 nM CYP2C9: 200 nM b5), 3 mM MgCl2, 20 µM BOMCC in 100 mM Tris-HCl buffer PH 7.4 and the reaction volume was 100 µL. The P450 reaction was started by addition of NADPH to a final concentration of 1 mM, and fluorescence was measured on a Spectramax M2e plate reader (Molecular Devices, Sunnyvale, CA, United States) at an excitation wavelength of 415 nm and an emission wavelength of 460 nm for BOMCC.

### 2.9 Assay of cytochrome P450 CYP2C19 Activity in Reconstituted Liposome System

The activity of CYP2C19 promoted by WT or mutant POR was tested using the fluorogenic substrate EOMCC (Invitrogen Corp, Carlsbad, CA, United States). The purified CYP2C19 (CYPEX, Dundee, Scotland, United Kingdom) was used to test the activities of the POR variants using 20 µM EOMCC as substrate. In vitro CYP2C19 assays were performed using a reconstituted liposome system consisting of WT/mutant POR, CYP2C9 and cytochrome b5 at a ratio of 5:1:1. The final assay mixture consisted of 2,5 µg DLPC (1,2-Dilauroyl-sn-glycero-3-phosphocholine) and proteins (0,5 µM POR: 100 nM CYP2C9: 100 nM b5), 3 mM MgCl2, 20 µM EOMCC in 100 mM Tris-HCl buffer PH 7.4 and the reaction volume was 100 µL. The P450 reaction was started by addition of NADPH to a final concentration of 0.5 mM, and fluorescence was measured on a Spectramax M2e plate reader (Molecular Devices, Sunnyvale, CA, United States) at an excitation wavelength of 415 nm and an emission wavelength of 460 nm for BOMCC.

### 2.10 Assay of cytochrome P450 CYP3A4 Activity in Reconstituted Liposome System

To compare the activities of CYP19A1 with another steroid binding cytochrome P450, we tested the effect of POR mutations to support the enzyme activity of CYP3A4. The activity of the major drug metabolizing enzyme CYP3A4 supported by WT or POR-284T was tested using the fluorogenic substrate**s** [BOMCC (7-Benzyloxy-4-trifluoromethylcoumarin) (Invitrogen Corp, Carlsbad, CA) as described earlier (Flück et al., 2010). The purified CYP3A4 (CYPEX, Dundee, Scotland, UK) was used to test the activities of the POR variants using 20 µM BOMCC. *In-vitro* CYP3A4 assays were performed using a reconstituted liposome system consisting of WT/mutant POR, CYP3A4 and cytochrome b_5_ at a ratio of 4:1:1 (POR:CYP3A4:b_5_). The final assay mixture consisted of liposomes and proteins (80 pmol POR: 20 pmol CYP3A4: 20 pmol b_5_), 2.5 mM MgCl_2_, 2.5 μM GSH and 20 μM BOMCC in 50 mM HEPES buffer and the reaction volume was 200 μl. The catalytic reaction was initiated by addition of NADPH to 1 mM final concentration, and fluorescence was monitored on a Spectramax M2e plate reader (Molecular Devices, Sunnyvale, CA) at an excitation wavelength of 415 nm and an emission wavelength of 460 nm.

### 2.11 Assay of cytochrome P450 CYP3A5 Activity in Reconstituted Liposome System

The activity of CYP3A5 promoted by WT or mutant POR was tested using the fluorogenic substrate BOMCC (Invitrogen Corp, Carlsbad, CA, United States). The purified CYP3A5 (CYPEX, Dundee, Scotland, United Kingdom) was used to test the activities of the POR variants using 20 µM BOMCC as substrate. In vitro CYP3A5 assays were performed using a reconstituted liposome system consisting of WT/mutant POR, CYP3A5 and cytochrome b5 at a ratio of 5:1:1. The final assay mixture consisted of 5 µg DLPC (1,2-Dilauroyl-sn-glycero-3-phosphocholine) and proteins (1 µM POR: 200 nM CYP2C9: 200 nM b5), 3 mM MgCl2, 20 µM BOMCC in 100 mM Tris-HCl buffer PH 7.4 and the reaction volume was 100 µL. The CYP3A5 reaction was started by addition of NADPH to 1 mM final concentration, and fluorescence was measured on a Spectramax M2e plate reader (Molecular Devices, Sunnyvale, CA, United States) at an excitation wavelength of 415 nm and an emission wavelength of 460 nm for BOMCC.

### 2.12 Statistical Analysis of results

Data are shown as mean, standard errors of the mean (SEM) in each group or replicates. Differences within the subsets of experiments were calculated using Student’s t-test. P values less than 0.05 were considered statistically significant.

## 3 Results

### 3.1 Effect of POR-P284T on Small molecule Reduction Activity

To study the effect of the P284T mutation on POR activity, we expressed WT and POR-P284T in *E.coli* and measured their ability to transfer electrons from NADPH to cytochrome c, MTT or ferricyanide. The POR-P284T mutation had a much lower capacity to reduce cytochrome c and MTT (**Table 1, Figure 2A and 2B**). Compared to WT POR, the P284T variant lost ∼50% of its activity to reduce cytochrome c and had only 15 % of the WT POR activity to reduce MTT. However, the ferricyanide reduction activity (**Table 1, Figure 2C**) was not affected by the P284T mutation. The loss of activities with cytochrome c and MTT indicates disruption of electron transport from NADPH to FAD, which could be due to conformational instability due to a P284T mutation affecting domain movements and transfer of electrons from NADPH to FAD.

**Table 1.** Kinetic parameters for activities of cytochrome c, MTT, ferricyanide reduction and CYP19A1 activity supported by POR-WT and POR-P284T variant. For the cytochrome c, MTT and Ferricyanide reduction assay, the NADPH concentration was fixed at 100 μM and varying concentrations of the substrate were used for the analysis. For the conversion of androstenedione to estrone, NADPH was fixed at 1 mM, and variable concentrations (10–1000 nM) of androstenedione were used. Data are shown as mean ± SEM of independent replicates.

|  | Km,<br>(µM) | Vmax<br>nmol/min/mg | Vmax/Km | % WT |
| --- | --- | --- | --- | --- |
| **Cytochrome c reduction assay** |  |  |  |  |
| WT | 4.6±0.7 | 128.2±6.1 | 27.6 | 100 |
| P284T | 3.7±0.8 | 48.7±3 | 13 | 47 |
| **MTT reduction assay** |  |  |  |  |
| WT | 19.9±1.8 | 179.5±4 | 9 | 100 |
| P284T | 48.3±7.5 | 64.8±3.1 | 1.3 | 15 |
| **Ferricyanide reduction assay** |  |  |  |  |
| WT | 18.9±2.4 | 600±19 | 31.8 | 100 |
| P284T | 6.9±1.6 | 248.1±11 | 36 | 113 |
|  | Km, Androstenedione<br>(nM) | Vmax<br>pmol/min/nmol | Vmax/Km | % WT |
| **CYP19A1; aromatase (androstenedione to estrone)** |  |  |  |  |
| WT | 80±14.5 | 0.72±0.03 | 0.009 | 100 |
| P284T | 165.7±147.9 | 0.12±0.037 | 0.0008 | 9 |

**Figure 2:**
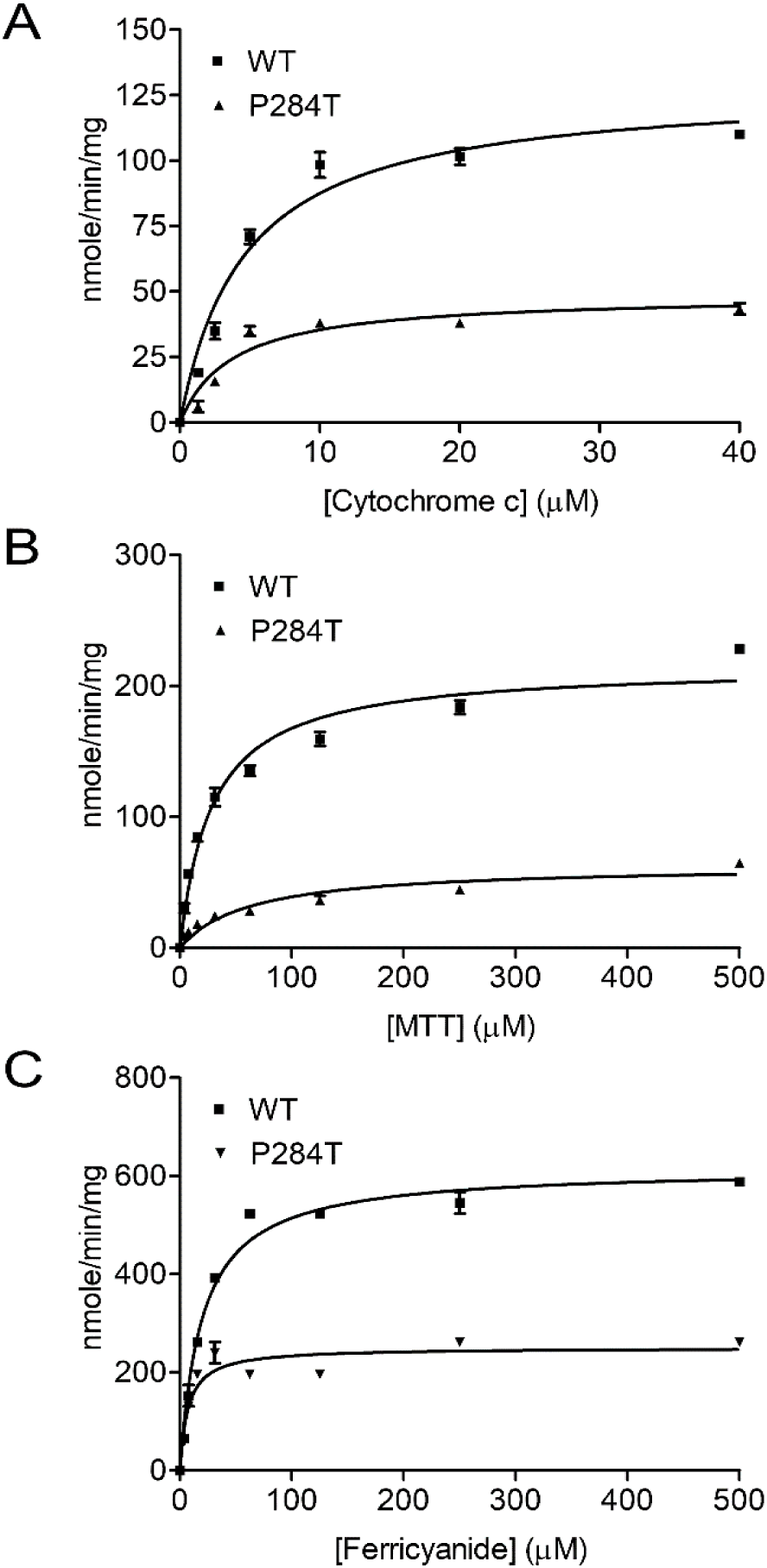
Cytochrome c, MTT and Ferricyanide reduction assay with WT and POR-P284T. (A) Cytochrome c, (B) MTT and (C) ferricyanide reduction assays were performed with the WT and POR-P284T variant. Kinetic assays were performed by monitoring the changes in absorbance at 550 nm for cytochrome c, 610 nm for MTT, and 420 nm for ferricyanide reduction. Data were fitted to the Michaelis-Menten kinetics model and analyzed using GraphPad Prism. The calculated Km and Vmax values are presented in Table 1.

### 3.2 Flavin Content

To differentiate the conformational changes and effects of POR mutation P284T on flavin binding, we evaluated the relative flavin content since the activity of POR may be affected by the changes in the binding of cofactors FMN and FAD. As compared to WT POR, both the FMN and the FAD-binding was affected due to P284T mutation. The FMN content of POR-P284T was 64% as compared to WT while the FAD content was reduced by 35% suggesting that POR-P284T affects both FMN as well as FAD binding to POR (**Figure 3**).

**Figure 3:**
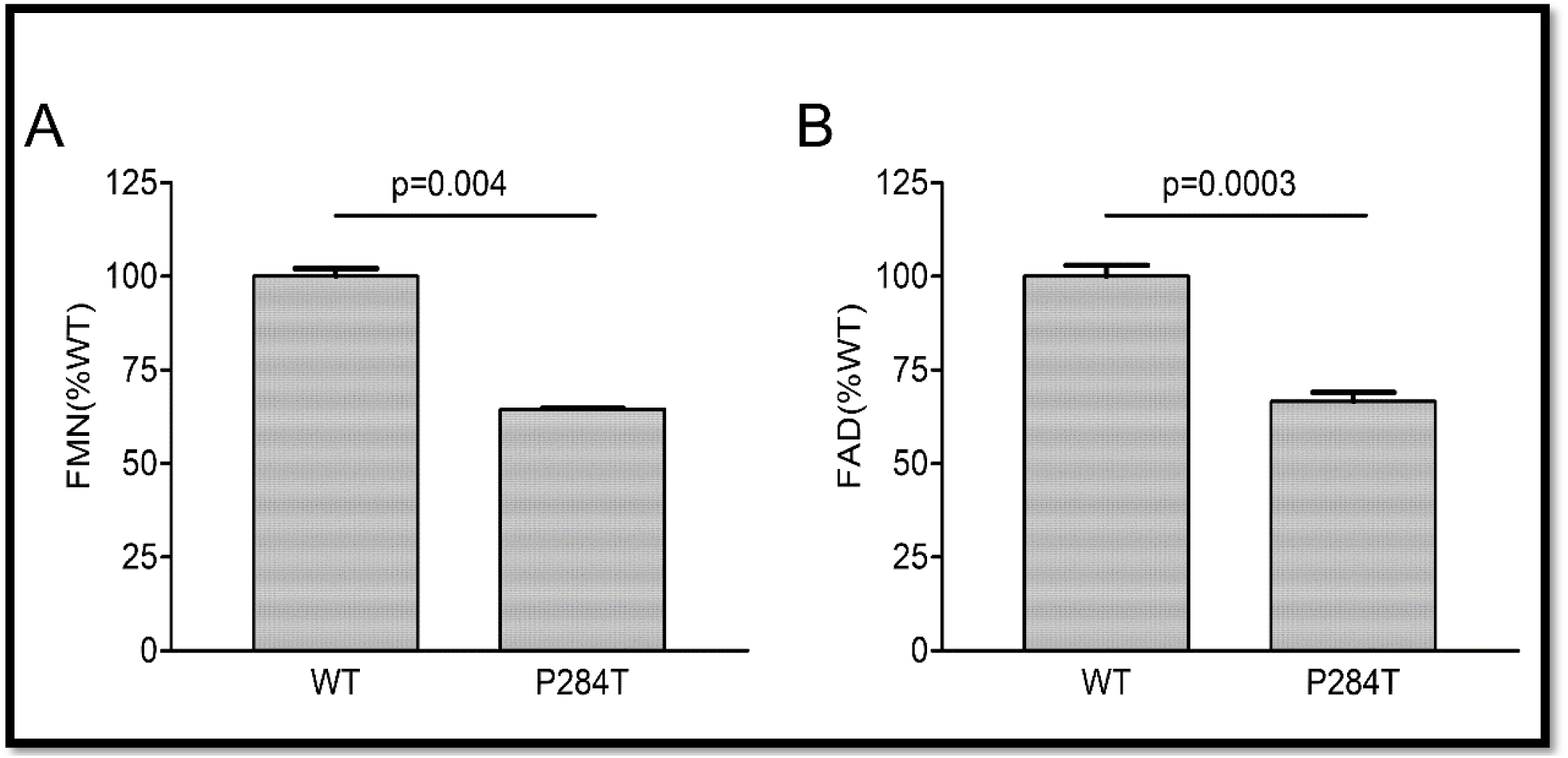
Flavin content of POR-WT and POR-P284T variant. Flavin content of the POR proteins was analyzed by boiling the proteins under selective pH conditions. Relative fluorescence unit (RFU) of the flavins released from the POR variants measured at (A) pH 7.7 (FMN or F_7.7_) and (B) pH 2.6 (FAD or F_2.6_) are shown. The RFU of WT POR was fixed as a hundred percent. Data are shown as mean ± SEM of three independent replicates.

### 3.3 CYP19A1-Aromatase Activity

The POR-P284T showed almost complete loss of CYP19A1 activity (**Figure 4, Table 1**). For the POR-P284T variant, the apparent Km for androstenedione was increased by 2-fold as compared to WT POR suggesting that P284T mutation affects either substrate interaction with CYP19A1 or the CYP19A1-POR interaction. The apparent Vmax of POR-P284T was reduced by ∼85%. The POR variant P284T showed only 9% residual activity in supporting CYP19A1 as compared to WT POR.

**Figure 4:**
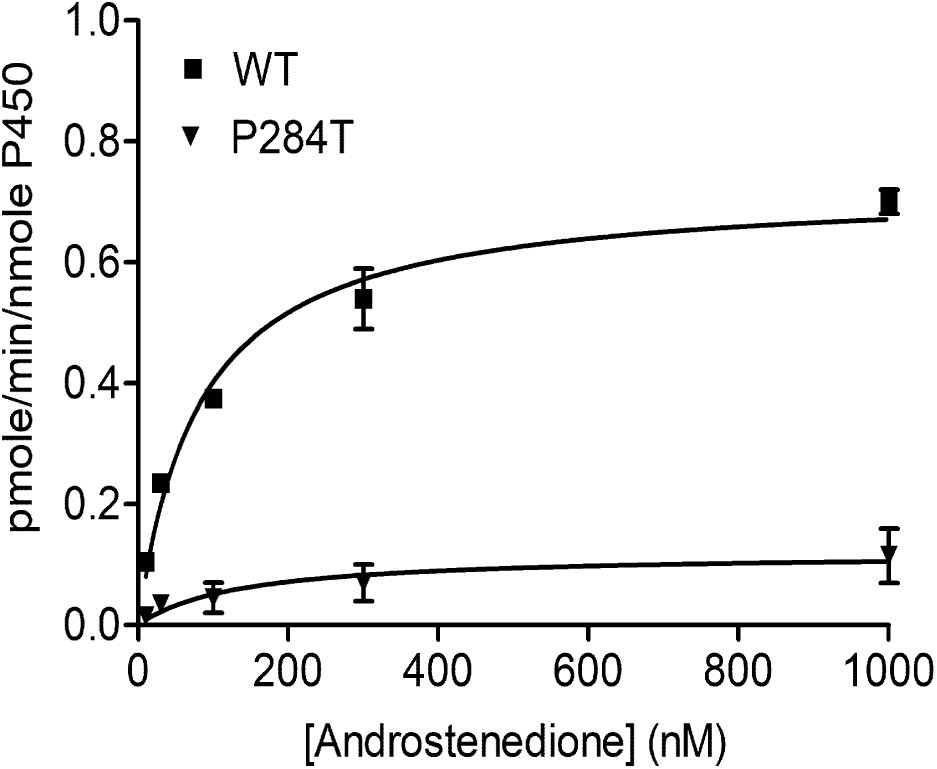
Enzymatic activity of CYP19A1 supported by POR-WT and POR-P284T variant. Bacterially expressed, purified, recombinant CYP19A1 and the enriched bacterial membranes containing POR proteins were inserted into liposomes and their activity to convert [^3^H] labeled androstenedione to estrone was tested by the tritiated water release assay. Data were analyzed using the Michaelis-Menten kinetics with GraphPad Prism. The calculated Km and Vmax values are shown in Table 1.

### 3.4 CYP2C9 Activity

We tested the activity of CYP2C9 supported by the WT and P284T variant of POR in reconstituted liposomes. Compared to WT POR activity of CYP2C19 supported by P284T variant of POR was reduced by 78% (**Figure 5**). This loss of more than three-quarters of activity compared to WT POR indicated a severe effect on drug metabolism supported by CYP2C9.

**Figure 5:**
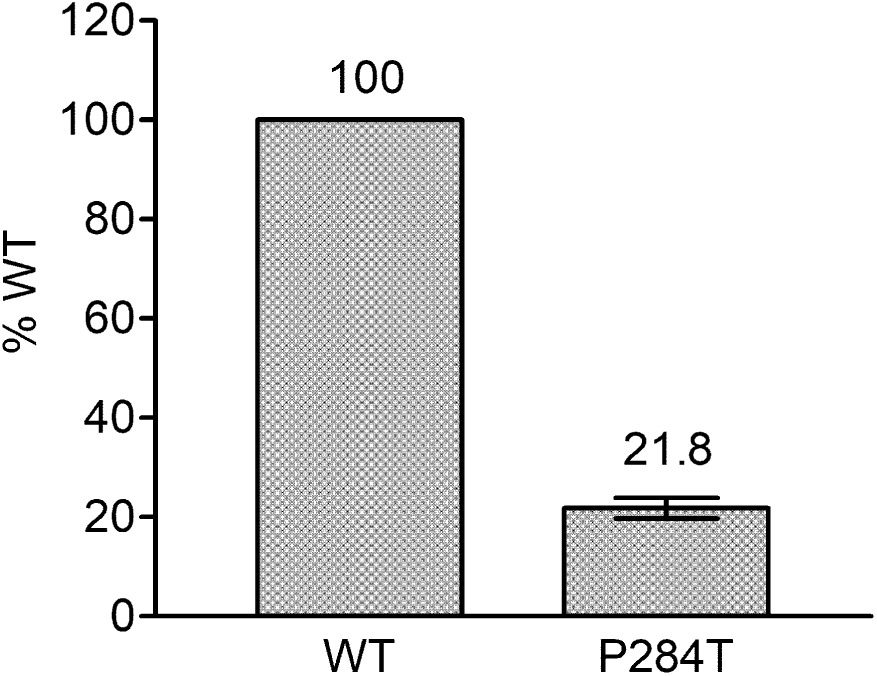
Activity of cytochrome P450 CYP2C9 supported by POR-WT and POR-P284T variant. Assay of CYP3A4 activity was performed to compare POR-WT and POR-P284T by using 20 µM BOMCC as a substrate. Activity with the WT POR was fixed as a hundred percent, and results are given as a percentage of WT activity. Data are shown as mean ± SEM of three independent replicates.

### 3.5 CYP2C19 Activity

The activity of CYP2C19 was tested with both the WT and P284T variant of POR. We found that in CYP2C19 assays the P284T variant of POR showed only 44% of the WT POR activity (**Figure 6**). The effect of P284T variation in POR on CYP2C19 was not as strong as in case of CYP2C9 activity, but the loss of activity was still more than 50%, indicating a reduced capacity of drug metabolism reactions mediated by CYP2C19.

**Figure 6:**
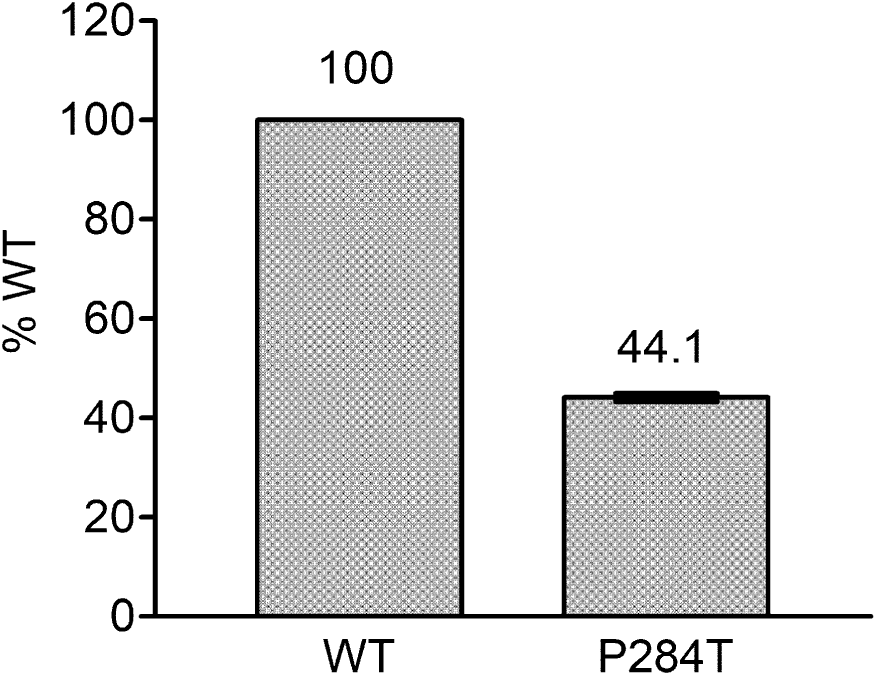
Activity of cytochrome P450 CYP2C19 supported by POR-WT and POR-P284T variant. Assay of CYP3A4 activity was performed to compare POR-WT and POR-P284T by using 20 µM EOMCC as a substrate. Activity with the WT POR was fixed as a hundred percent, and results are given as a percentage of WT activity. Data are shown as mean ± SEM of three independent replicates

### 3.6 CYP3A4 Activity

The P284T variant of POR showed only 22.8 % activity as compared to the WT enzyme (**Figure 7**). The loss of activities for drug metabolizing cytochrome P450 enzyme-CYP3A4 by the POR variant P284T indicates problems with POR-P450 interactions which seem to be different for different cytochrome P450 partners.

**Figure 7:**
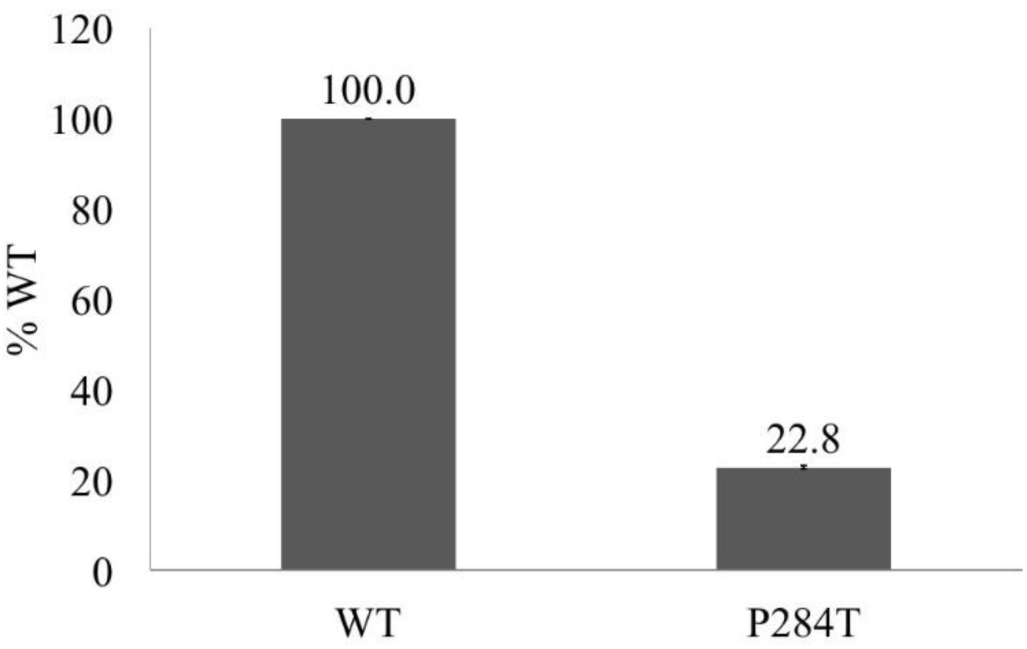
Activity of cytochrome P450 CYP3A4 supported by POR-WT and POR-P284T variant. Assay of CYP3A4 activity was performed to compare POR-WT and POR-P284T by using 20 µM BOMCC as a substrate. Activity with the WT POR was fixed as a hundred percent, and results are given as a percentage of WT activity. Data are shown as mean ± SEM of three independent replicates.

### 3.7 CYP3A5 Activity

The 284T variant of POR had only 43.8% of the WT activity in CYP3A5 assay (**Figure 8**). This was different from the results obtained for the CYP3A4 activity assays, indicating there are differences in the interaction of POR with these two closely related cytochrome P450 proteins.

**Figure 8:**
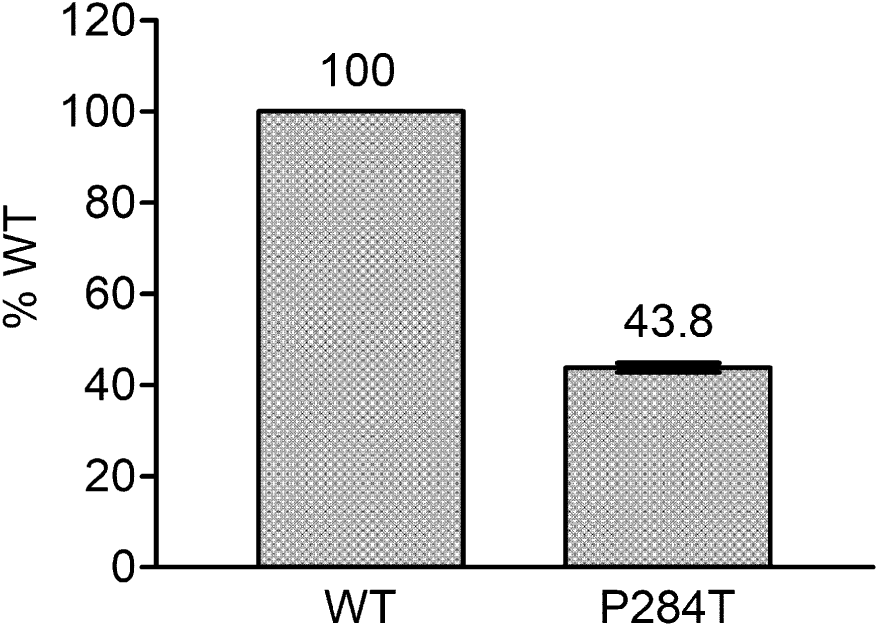
Activity of cytochrome P450 CYP3A5 supported by POR-WT and POR-P284T variant. Assay of CYP3A5 activity was performed to compare POR-WT and POR-P284T by using BOMCC as a substrate. Activity with the WT POR was fixed as a hundred percent, and results are given as a percentage of WT activity. Data are shown as mean ± SEM of three independent replicates.

## 4 Discussion

In previous reports, the effects of POR variants on drug metabolizing cytochrome P450 enzymes has not been studied in detail, and effects on most major drug metabolizing enzyme activities remain unknown. The cytochrome P450 subfamily 3A is the most versatile among all P450 enzymes and accounts for the metabolism of a large number of xenobiotics and pharmaceutical compounds. Cytochrome P450 3A4 (CYP3A4) is the principal hepatic enzyme that metabolizes a large percentage of drugs and endogenous substrates (Klein and Zanger, 2013;Meyer et al., 2013;Zanger and Schwab, 2013). While CYP3A5 also has a significant role in xenobiotic metabolism in extrahepatic tissues like intestine and gut. The cytochrome P450 subfamily 2C is the second most relevant class of P450 proteins, and CYP2C9 and CYP2C19 are major drug-metabolizing enzymes.

Our result shows that mutations in POR can affect different activities of POR to varying extents and some cytochrome P450 proteins are affected more than others. Also, there may be differences due to different allelic variations of cytochrome P450 proteins. The effects of different cytochrome P450 isoforms on the impact of POR activities has not been studied in detail. One study (Subramanian et al., 2012) tested different alleles of CYP2C9 (*CYP2C9.1, CYP2C9.2,* and *CYP2C9.3*) with a small number of POR variants but patterns were similar for all isoforms of CYP2C9 (Subramanian et al., 2012).

POR mutations leading to the significant loss of FAD/FMN co-factors hurt all redox partner activities. However, in most other cases, the effects of POR mutations on activities of the individual redox partner could be variable. For the mutation P284T, we observed a small loss of cofactor binding, which could be due to overall protein stability changes due to the location of the P284 residue near the hinge region of POR that joins FMN binding domain with the FAD-binding domain. However, the activity of ferricyanide reduction which represents direct electron transport through FAD without the involvement of FMN indicates that this loss of FAD did not severely reduce the overall quality of the protein. Cytochrome c reduction that involves protein-protein interaction and MTT reduction which a dual electron transfer system through FMN showed loss of activities.

The location of the P284T mutation in the hinge region indicated a potential impact on domain movements which may change protein-protein interaction which may differ based on which redox partner is being tested. To test this hypothesis we tested POR-P284T with several different cytochrome P450 proteins. The CYP19A1 activity requires the transfer of three pairs of electrons, and therefore, is more susceptible to change in protein interaction with POR (Pandey et al., 2007;Flück and Pandey, 2017). A severe loss of CYP19A1 activity due to P284T mutation (9% of WT activity) indicated a significant effect on interaction with POR. In case of drug metabolizing cytochrome P450s, CYP2C9 and CYP3A4 lost more than 75% of activities while CYP2C19 and CYP3A5 lost more than 50% of the WT POR activity due to P284T mutation in POR. Taken together these results indicate variability in interaction with different cytochrome P450s due to P284T mutation in POR. These results further underscore the fact that effects of POR mutations can be quite variable for different redox partners and mutations in POR require detailed characterization with different partner proteins and small molecule substrates to understand the biochemical consequences of individual POR mutations.

Therefore, biochemical analysis using recombinant proteins is necessary to confirm the damaging effects of mutations with each redox partner separately (Parween et al., 2016;Udhane et al., 2017;Flück and Pandey, 2019). Several cytochrome P450 proteins for which POR is the redox partner, still have uncharacterized activities and their physiological roles are not defined. These, other partners of POR could also influence the metabolic profiles of patients with PORD. Preliminary computational docking and functional analysis suggest that the loss of activity caused by some POR variants may be reversed by the introduction of external FMN (Nicolo et al., 2010) or FAD (Marohnic et al., 2006). However, the effect of flavin treatment in PORD needs to be tested in a clinical setting.

PORD is a complex disorder affecting both the steroid and the drug metabolism in the affected individual with varying effects. Different POR variations can cause highly variable effects on different metabolic reactions depending on the unique nature of the interaction of POR with different redox partners. A careful examination of affected target proteins by use of urine/serum steroid analysis is required for initial verification of PORD. In cases of severe loss of drug metabolizing cytochrome P450 enzyme activities, further considerations about the clearance of drugs in patients with PORD would require evaluation. Effect of altered clearance of kidney transplant drug tacrolimus has been indicated in several studies among the people with POR*28 allele (Zhang et al., 2015). Effects of different POR variations found in non-clinical populations should also be investigated in detail, and our recent work has shown that some of these variations in POR could lead to severe changes in activities of enzymes dependent on POR (Udhane et al., 2017).

## 6 Author Contributions

Participated in research design: AP

Conducted experiments: SP, MV, SU

Contributed new reagents or analytical tools: NK

Performed data analysis: SP, MV, AP

Overall supervision of the project: AP

Wrote or contributed the writing of the manuscript: SP, MV, SU, NK, AP

## 8 Acknowledgments

Prof. Walter L. Miller (UCSF, San Francisco, CA, USA) provided several POR cDNA clones.

